# Unveiling the Secrets of Extracellular Vesicles in Urban Water Systems: Understanding the Link Between Human and Environmental Health

**DOI:** 10.1101/2024.05.15.594346

**Authors:** Fei Liu, Yi Li, Yunxian Piao, Yong Wang, Zaiyao Liang, Luke P. Lee

**Author notes:** Corresponding author (F. Liu); (L. P. Lee).

## Abstract

It is crucial to gain valuable insights into the ecological health of rivers to inform management decisions and protect sustainable living conditions. Understanding biological information is vital to gaining insight into river ecosystem biodiversity, but reliable methods are challenging. Here, We investigate the potential impact of extracellular vesicles (EVs) in urban water systems on human and environmental health to promote urban sustainability. We used EXODUS to detect EVs and perform metaproteomic analysis on samples from an urban water system that contained human feces, wastewater, bacteria, plants, arthropods, and soil. We analyzed EVs collected from urban and green areas, observing taxonomic variations and discovering bacterial contributions to their protein content. According to our research, the abundance and expression levels of proteins in EVs can indicate how human activities affect microbial communities in rivers, potentially impacting public health. Our study offers a fresh perspective on the interconnectedness of urban sustainability, public health, and river ecosystem biodiversity.

## Introduction

Rapid urbanization in densely populated areas has presented significant environmental challenges for urban rivers and sustainability. Despite serving multiple functions such as local climate regulation, biodiversity preservation, and provision of water resources^1^, urban expansion has had a detrimental impact on the health of these rivers^1^. Human activities can cause damage to natural habitats and water quality, leading to a decline in biodiversity and the essential ecosystem services they provide. The alarming decline of a particular species has dire implications that can negatively impact biodiversity conservation and public health^2-5^. Urban areas play a crucial role in enhancing human well-being through services such as air purification, noise reduction, cooling effects, and runoff management^6,7^. Urban river ecosystems that are left undisturbed can sustain various wildlife populations within city limits^8^. Urban biodiversity is the variety of living organisms in an urban area, including plants, animals, and bacteria^9,10^. While current assessments primarily focus on broad categories like ecosystem types or expanses alongside species diversity and abundance^9^, this raises the question: How can we develop strategies and methods to improve the precision analysis at the molecular level and the monitoring of biodiversity affected by human activities?

Organisms release membrane-enclosed structures called EVs that carry molecular signals, such as DNAs, RNAs, proteins, lipids, and metabolites, into the environment^11-13^and harbor valuable information about the biological components of an ecosystem^14-18^. In urban rivers, microorganisms exchange genetic material and molecular information via EVs^19^, aiding diverse species to adapt and survive challenging environments^20-23^. The transfer of EV cargo can increase the genetic diversity of microorganisms^24,25^, which may enhance their ability to withstand pollutants and other stressors commonly encountered in urban waterways^26^. Early studies have demonstrated that *Prochlorococcus*, a prevalent marine cyanobacterium, releases EVs containing proteins, DNA, and RNA. Coastal and open-ocean seawater samples also exhibit diverse bacterial DNA-carrying EVs^24^. *Prochlorococcus* EVs support the growth of other bacteria and play a significant role in marine carbon flow. It highlights the contribution of EVs to information exchange within marine microbial communities by transporting various nutrients and signaling molecules, adding to community complexity^27^. However, this research primarily. It focuses on laboratory cultures of *Prochlorococcus* and may need to comprehensively capture the intricacy and diversity of natural EV production in marine ecosystems^28^. Other studies reveal variations in EV production among different strains of cultivated marine microbes and their correlation with critical environmental variables^29^. This study establishes a quantitative framework for understanding the role of EVs in marine ecosystems and enhancing our ecological and biogeochemical knowledge of the ocean^29^. However, the paper mainly investigates the variations in EV production and size, with limited exploration of the specific environmental functions played by these EVs within the marine ecosystem; therefore, further research is necessary.

In this study, we report on how to uncover the hidden treasure of EVs in urban water systems to discover the interconnectedness of urban sustainability, public health, and river ecosystem biodiversity. We utilized EXODUS^30^ to detect EVs and conduct metaproteomic analysis on samples from an urban water system that contained human feces, wastewater, bacteria, plants, and soil. We analyzed EVs collected from urban and green areas, observed taxonomic variations, and discovered bacterial contributions to their protein content.

First, we collected EVs from urban sewer water, water body bacteria, and green spaces. These samples were collected at six different locations along the Guanlan River, an urban river in Shenzhen, China. We then comprehensively analyzed the protein expression levels in the enriched EVs isolated from these water samples. (Figure 1a). The Guanlan River, located centrally within Shenzhen, encompasses diverse historical, geographical, and topographical features that make it an exemplary urban river (Figure 1a S1 - S3, S5). Its catchment area includes urban community areas, schools, hospitals, shopping malls (Figure 1a S1 - S3, S5), green spaces (Figure 1a S4, S6), agricultural lands, and natural reserve zones.

**Figure 1.**
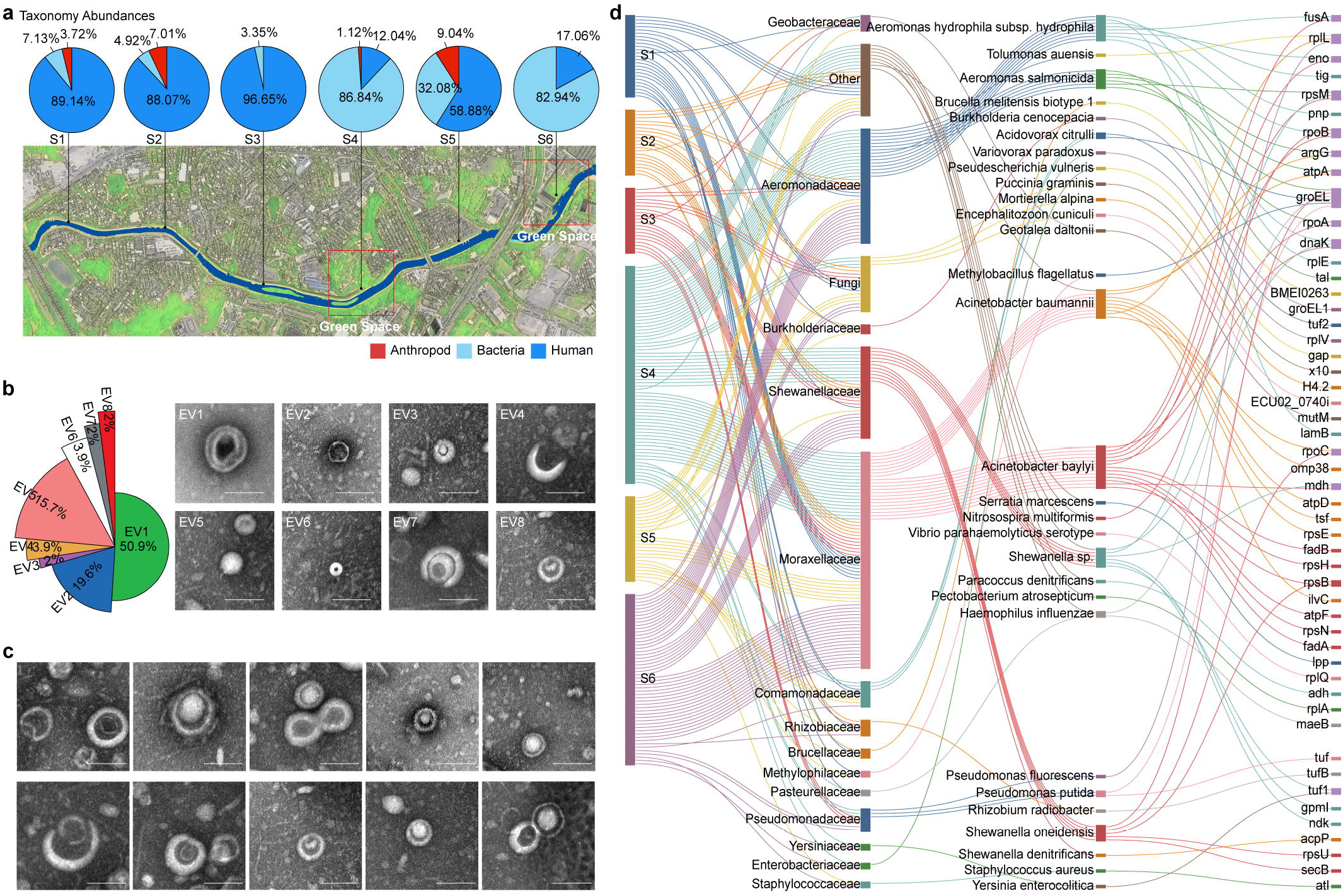
Environmental EV analysis with molecular-level resolution depicts the microbe taxonomic group change from different sample collection positions. **a**. The samples were collected from specific points along the Guanlan River, and within each sample, the predominant proteins originated primarily from three primary sources: arthropods, bacteria, and humans. These samples undergo comprehensive analysis to identify the main protein categories present, including visual representations such as typical TEM images showcasing the characteristic appearance of these proteins (scale bar: 200nm). **b**. When observed through TEM, determining the proportion or ratio of EVs falls into distinct morphological categories. This involves analyzing the distribution or relative abundance of EVs based on their specific structural characteristics as identified or classified by TEM imaging (scale bar: 100nm). **c**. TEM unveiled a multilayer from multiple samples under TEM. Most multilayer EVs have two layers of the membrane with different thicknesses. **d**. Differentially abundant in taxonomic groups and associated genes are shown in the Sankey diagram. a. scale bar: 200nm, b. scale bar: 100nm.

Consequently, the EVs found in this river may carry signatures of urban pollution and reflect agricultural activities and natural environments. Given the varied topography comprising estuaries, wetlands, river channels, and riverbanks, the distribution and composition of EVs are likely to differ across these distinct habitats, making the Guanlan River an ideal subject for comprehensive exploration into their dynamics and functions within this ecosystem. Investigating these dynamics holds excellent promise for gaining insights into the intricate workings of this river’s ecosystem.

Our investigation has three main research questions that we aim to address. These include: 1. What are the metaproteomic characteristics of urban rivers? 2. How does the expression level of EV proteins correlate with biodiversity in urban rivers? 3. What are the reasons behind changes in biodiversity of urban rivers due to human activities, and how do these changes subsequently impact human health? Based on our research, the quantity and quality of proteins found in EVs can indicate how human activities impact microbial communities in rivers, affecting public health. Our research offers an innovative viewpoint on the relationship between urban sustainability, public health, and the biodiversity of river ecosystems. We found that the abundance and expression levels of proteins in EVs can indicate how human activities affect river microbial communities. This can impact public health by altering the river ecosystem’s biodiversity.

## Results

The peptides generated through enzymatic digestion of proteins extracted from EVs at six different sampling locations along the Guanlan River were used for the metaproteomic analysis using tandem mass spectra (MS/MS). Most of each sample comprised three primary sources: arthropods, bacteria, and humans (Figure 1a). These various sources contributed collectively to the abundance and diversity of proteins within the urban river ecosystem.

A diverse range of EVs, with various sizes and shapes, indicates the existence of specialized subpopulations with distinct functions characterized by lipid bilayers and internal structures. We employ Transmission Electron Microscopy (TEM) to analyze EVs with multiple structures, which provides accurate size measurements and useful morphological information (refer to Figure 1b). Our findings show that EVs from human-derived samples primarily consist of single structures, whereas EVs from bacteria have a higher incidence of double-EV structures. Further investigation classified these EVs into eight categories (Figure 1b), with single (Figure 1b. EV1, EV2, EV4, EV5, EV6), double (Figure 1b. EV3, EV8), and small double EVs being the most prevalent (Figure 1b. EV7). Additionally, we discovered the ubiquitous presence of double-layered and multiple-layered extracellular EVs in the urban river (Figure 1c). With the increasing volume of TEM data (Figure S2 - S14), Manual analysis of TEM images is time-consuming and labor-intensive, requiring skilled experts to interpret complex information regarding EV structure. In this study, we employed the open-source Vision Transformer (ViT) model^31^ for the automated classification of TEM images to facilitate our analysis (Figure S25). AI’s scalability allows efficient handling of extensive datasets, supporting objective interpretation and comprehensive exploration of diverse EV nanostructures. Specifically, Artificial Intelligence in EV Morphology Classification (AIVMC) has been applied to classify EV TEM images.

The Sankey diagram in Figure 1d shows the flow of microbiome information between different taxonomic groups. The width of each node corresponds to the relative abundance of proteins in samples collected from various locations. In contrast, the thickness of the connections between nodes indicates the quantity of microbial taxa transitioning between groups. Taxonomic categories, ranging from bacterial families to individual genes, are organized hierarchically, facilitating a comprehensive understanding of microbial population distribution and evolution within the urban river ecosystem. This graphical representation yields valuable insights into dynamics within microbial communities: green spaces exhibit a prevalence of aerobic bacterial communities, whereas densely populated areas demonstrate a decrease in total protein content with anaerobic and facultative anaerobic bacterial populations predominating. Additionally, metaproteomic analysis highlights the distribution and transfer of microbial diversity across various taxonomic levels.

We analyze and visually represent the distribution and variability of total proteins within each sample group. The concise visualization of metaproteomic data across diverse samples in Figure 2a reveals the distribution and variability of protein abundance within each sample. The varying heights of the boxes, particularly taller in samples S1, S2, S3, and S5 and shorter in samples S4 and S6, indicate distinct levels of variability. Whiskers extending from the boxes represent normalized protein expression levels beyond the interquartile range (IQR), while outliers are depicted as individual points. This graphical representation of the discrepancy of protein abundance distributions between green spaces and densely populated areas highlights variations in both variability and central tendency.

**Figure 2.**
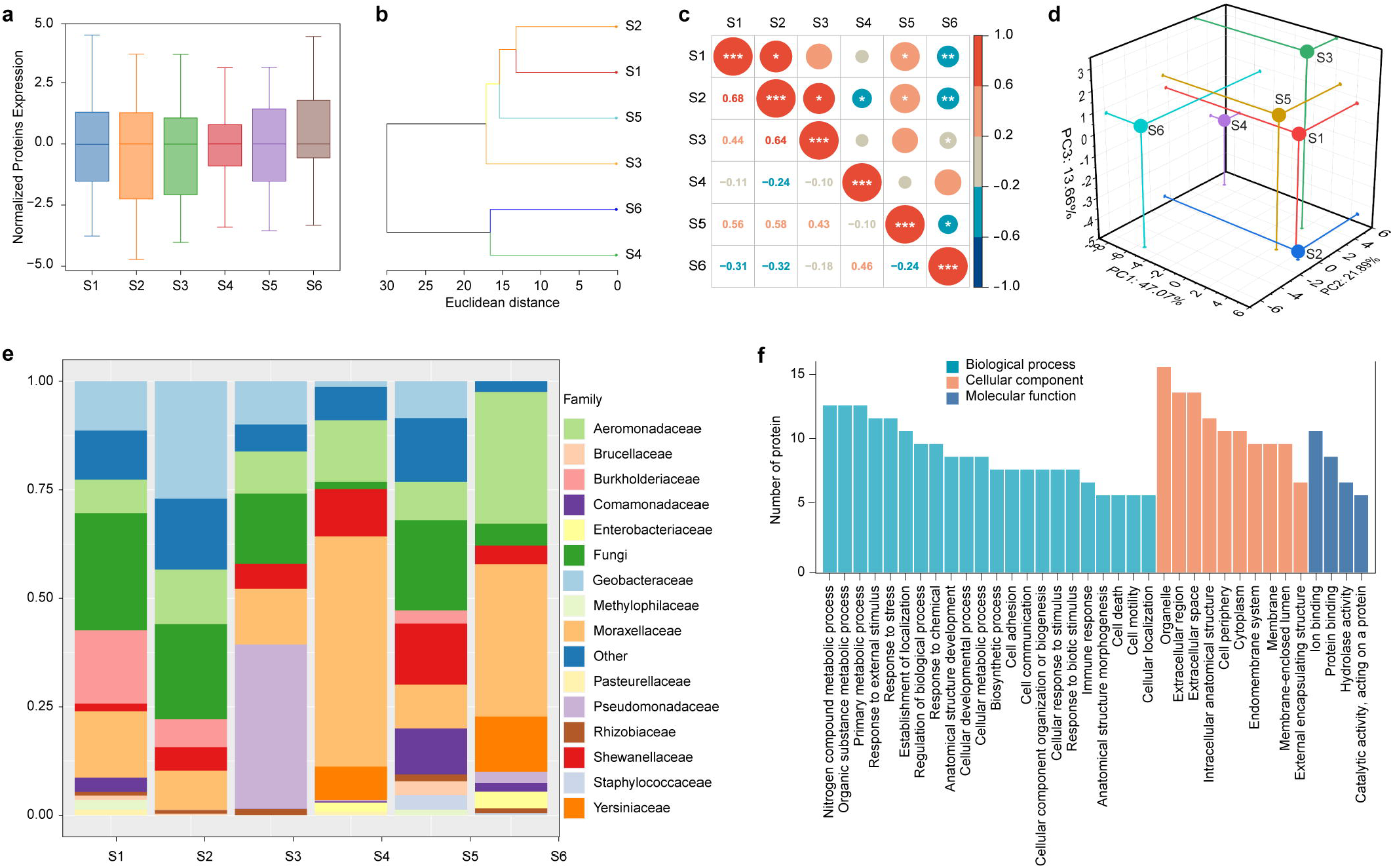
Categorized water samples highlighted a prevalence of bacteria significantly influenced by urban river ecosystems. **a**. Normalized protein expression levels and accurate quantitative comparisons of protein abundance across different samples corrected the sample-to-sample and technical variations. **b**. The unsupervised hierarchical clustering technique sorted the six water samples into two clearly defined groups based on their inherent similarities or differences. **c**. The proteomic profile correlation matrix of different water samples highlighted resemblances in the proteomic data across various water samples. Specifically, samples S4 and S6 displayed a direct, positive correlation in their protein profiles, while S1, S2, S3, and S5 also demonstrated a similar positive relationship among themselves. However, what stood out was that S4 and S6 showcased a contrasting negative correlation with respect to S1, S2, S3, and S5, suggesting notable distinctions in their overall protein compositions or profiles. d. Identifying clusters of water EV samples using 3D PCA illustrates relationships among samples across different dimensions. Each plotted point represents a replicate from a specific experimental group distinguished by unique colors for clarity. **e**. Analysis of microbial community abundance reveals that fungi are more abundant in S1, S2, S3, and S5, while bacterial communities are more abundant in S4 and S6. The decrease in water oxygen levels caused by human feces can result in an increase in fungal proliferation, which significantly impacts the biodiversity of urban rivers. **f**. The top groups of GO terms with the highest number of representative proteins were related to nitrogen compound metabolism, organic substance metabolism, and primary metabolic processes. These categories were significantly upregulated in the bacterial community, highlighting the crucial role of bacterial communities in the nitrogen cycle of urban rivers and emphasizing that processes related to bacteria form the foundation of diversity in urban river ecosystems diversity.

The water samples comprised a diverse array of EVs, encompassing single EVs of varying sizes. Here, we employed the unsupervised hierarchical clustering method; the six water sample sets were effectively classified into two distinct clusters, as depicted in Figure 2b. Hierarchical clustering enables the identification of coherent groupings and unveils hidden patterns. For instance, samples S1, S2, S3, and S5 merged into a cohesive cluster, while samples S4 and S6 formed a separate cluster based on their collective protein abundance and Euclidean distances.

We further elucidate associations and dependencies within the samples through correlation analysis, revealing similarities in proteomic values among different water samples (Figure 2c). Samples S4 and S6 exhibit a positive linear relationship, while S1, S2, S3, and S5 also display such a correlation. However, the four samples exhibit a unique negative linear relationship with S4 and S6. These samples and S4 and S6 have a distinct negative linear relationship, indicating significant differences in their overall protein profiles.

We illustrate the interrelationships among the samples along three orthogonal principal axes by utilizing three-dimensional principal component analysis (3D - PCA). To present the protein expression data in Figure 2d. Here, PCA offers distinct benefits in metaproteomic analyses by reducing dimensionality into a manageable format while retaining essential information within the samples. Each point on the plot represents a replicate from an individual experimental group, color-coded for differentiation. By incorporating all quantified proteins, PCA effectively segregates the samples into two distinct clusters along orthogonal principal components (PC1 - PC3), accounting for 47.07%, 21.89%, and 13.66% of the total variance in proteomic data. Each principal component represents a different direction in the feature space of the data and explains a certain percentage of the total variance. Samples S1, S2, S3, and S5 exhibit proximity while demonstrating noticeable separation from samples S4 and S6, indicating a significant disparity between these groups.

Conversely, the observed shift in bacteria communities in S4 and S6 suggests an increase in oxygen levels in regions dominated by aerobic bacteria such as *Moraxellaceae, Aeromonadaceae, Shewanellaceae*, and *Yersiniaceae* (Figure 2e, Table S3). This phenomenon will likely promote biodiversity development in urban rivers that typically improve water quality. Increased biodiversity is crucial in regulating water pollutants, contributing to cleaner and safer water sources. Consequently, this directly impacts the provision of urban populations with access to clean drinking water. Moreover, urban rivers teeming with diverse species can offer recreational opportunities like fishing, boating, and hiking. Access to green spaces and natural areas has been scientifically proven to positively affect mental health and well-being by reducing stress and promoting relaxation^32,33^. The microbial community composition depicted in Figure 1e demonstrates a predominance of fungi in samples S1, S2, S3, and S5, while bacterial communities dominate in samples S4 and S6. As human activities decrease within these investigated areas, a noticeable transition from fungal populations to certain anaerobic bacteria families such as *Burkholderiaceae, Moraxellaceae, Geobacteraceae, Pseudomonadaceae*, and *Shewanellaceae* occurs towards aerobic or facultative anaerobic bacteria including *Moraxellaceae, Aeromonadaceae, Shewanellaceae*, and *Yersiniaceae*. The shift primarily reflects significant changes in water oxygen levels influencing dominant bacterial communities. The decline in water oxygen levels and fungal overgrowth caused by human fecal contamination can profoundly impact urban river biodiversity. Reduced oxygen levels threaten aquatic organisms, leading to decreased species diversity. Additionally, fungal overgrowth alters microbial community compositions, affecting overall ecosystem dynamics.

Our gene ontology (GO) analysis revealed prevalent categories linked with nitrogen compound metabolism, organic substance metabolism, and primary metabolic processes across all samples (Figure 2f). These interconnected biological pathways are closely associated with microbiome functions. The enhanced expression of metabolic activities within the bacterial community offers valuable insights into encapsulated EVs harboring urban river microbiomes’ functional potential. Particularly noteworthy is the significant upregulation observed in crucial processes, including nitrogen compound metabolism, organic substance metabolism, and primary metabolic pathways, emphasizing the fundamental role played by bacterial communities in driving the nitrogen cycle dynamics within urban rivers. This highlights that microbial-driven mechanisms are a cornerstone for maintaining diversity within urban river ecosystems.

The Sankey diagram illustrates the connection between upregulated genes in humans and the relative abundance of GO items, which subsequently flow into the three major GO domains (Figure 3a). Our research shows a higher number of proteins from human organs such as the kidney, placenta, ovary, and skin in the water samples taken from urban rivers. This indicates that the leading cause of pollution in these rivers is most likely due to contamination from human feces or sewage (Figure 3b).

**Figure 3.**
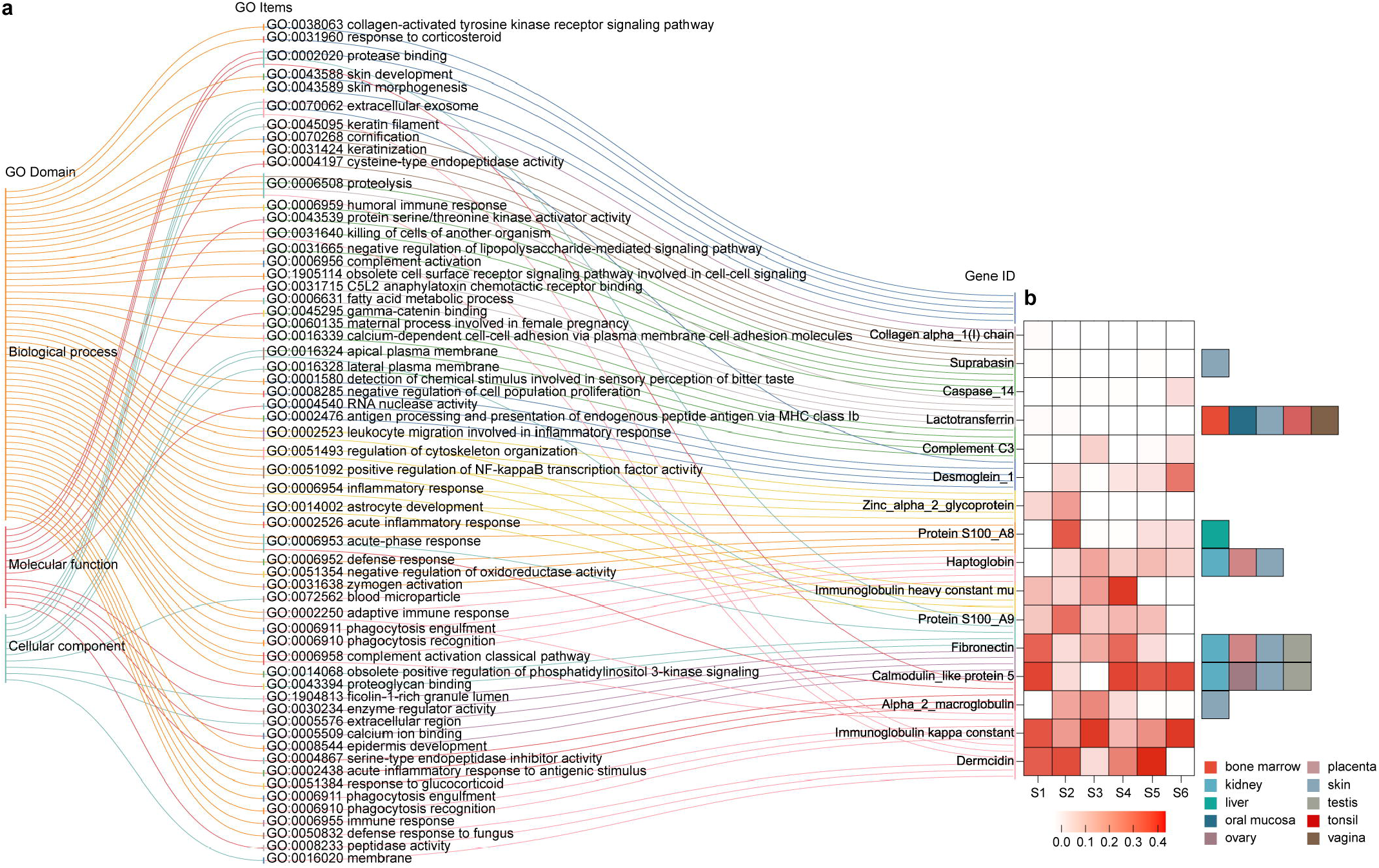
The protein abundance, expression level, and alternation involvement in pathways in EVs reflect transformations in microbial communities brought about by human activities, with potential implications for human health. **a**. The Sankey diagram represents relationships between GO items identified by their gene IDs within specific GO domains. **b**. Heatmap showing the relative abundance of identified EV proteins from human sources among subgroups. The water sample contained a relatively higher number of proteins from human sources, specifically originating from the kidney, placenta, ovary, and skin. This indicates that human feces or sewer water is the primary source of contamination in urban river samples.

Concurrently, the coexistence of human fecal matter, potential stormwater runoff carrying bacterial and sewer water overflow can trigger algal blooms due to an excess of nutrients^34^, such as those from fertilizer, wastewater, and stormwater runoff, in conjunction with optimal conditions like abundant sunlight and warm temperatures. Algae blooms can proliferate rapidly, impacting water quality by altering clarity and oxygen levels. Monitoring is necessary to assess the potential for harmful toxins produced by cyanobacteria, a type of algae^35-37^. Frequent or prolonged algal blooms are signs of stress within water bodies, which can have adverse effects such as nutrient enrichment, habitat degradation, and reduced support for aquatic life^38,39^. *Cyanobacteria* blooms, in particular, can produce dangerous toxins that can cause illness or death in humans and animals and create dead zones within the water system^40-42^.

Furthermore, they escalate treatment costs for drinking water facilities and pose challenges for industries reliant on clean water resources^5,43^. These alterations in biodiversity disrupt the natural equilibrium of rivers, resulting in issues related to water quality management, disease vectors, and other ecological imbalances that indirectly impact human health through the river’s role as a source of freshwater supply that supports local ecosystems^44^. Ultimately, biodiversity modification within urban rivers has intricate implications for human health and well-being, encompassing direct health hazards and broader impacts on overall quality of life^45,46^.

## Discussion

The findings emphasize urban rivers’ crucial role, highlighting their multifaceted significance in climate regulation, biodiversity support, and water provision for urban development. EV analysis is a high-resolution and cost-effective methodology that shows promise for comprehensively investigating urban river ecosystems and biodiversity. It provides an all-encompassing perspective on these ecosystems, including visible species, microorganisms, and the food web – essential for effective ecosystem management. Our findings can contribute to understanding urban river ecosystem biodiversity and health through EV analysis, thereby informing conservation efforts. This knowledge can guide policymakers and conservationists in protecting and restoring these ecosystems. EV analysis is a robust tool for continuous environmental monitoring that enables early detection of pollution and invasive species while facilitating prompt and effective pollution control measures.

In conclusion, our report is critical in evaluating the impact of urban river health on human well-being and activities. Understanding the complex connections between urban river health, pollution, and public health is crucial to uncovering the hidden treasures of EVs from urban water systems. This will provide valuable insights for targeted interventions and educational campaigns to improve the well-being of communities living near these rivers. In conclusion, our report is vital in enhancing community well-being by shedding light on the intricate relationships between urban river health, pollution, and public health.

## Methods

### EV isolation

After a 1 μm prefiltration step, the water sample was isolated using the EXODUS method with an alternative vacuum actuation at −20 kPa and a conversion time of 10 seconds. Subsequently, each subfraction was eluted twice with PBS via EXODUS. Finally, the three reserved fractions were recovered in 200 μl of 1X PBS and stored at −80°C for subsequent analysis.

### Transmission electron microscopy (TEM)

The EV samples were combined with a 4% paraformaldehyde solution in a 1:1 ratio and incubated on Formvar carbon-coated grids for 20 minutes, followed by a wash with PBS. Subsequently, the EVs were fixed with 1% glutaraldehyde for 5 minutes, rinsed with distilled water, and then exposed to a 2% uranyl acetate solution at room temperature for 30 seconds. After air-drying, the samples were observed using a transmission electron microscope (FEI Talos F200) operating at 80 kV.

### Quantitative proteomic analysis

The analyses were conducted using a Q-Exactive HF mass spectrometer (Thermo, USA) equipped with a Nanospray Flex source (Thermo, USA). Samples were loaded and separated on an EASY-nLCTM 1200 system (Thermo, USA) with a C18 column (25 cm × 75 μm). The flow rate was set at 300nL/min and a linear gradient of 90 min was applied (0∼1 min, 0%-2% B; 1∼2 min, 2%-6% B; 2∼51 min, 6%-21% B; 51∼70 min, 21%-31% B; 70∼81 min, 31%-43% B; 81∼84 min, 43%-100% B; 84∼90 min, 100%B; mobile phase A = 0.1% FA in water and B = 0.1% FA in 80% CAN and 19.9% water). Full MS scans were acquired within the mass range of 350-1650 m/z at a resolution of 60000 with an AGC target value set at 3e6. The top 20 most intense peaks in the MS spectra were subjected to higher-energy collisional dissociation fragmentation using HCD with a collision energy of 28. MS/MS spectra were obtained at a resolution of 30000 with an AGC target value set to 2e5 and a maximum injection time limited to 80 ms. The Q Exactive HF dynamic exclusion was set for 40.0 s under positive mode.

### Database searching of MS data

The LC-MS/MS raw data were imported into Maxquant (Version 1.6.17.0) for label-free quantification analysis utilizing the Andromeda search engine. A target-decoy-based approach was employed to control for false discovery rate (FDR) and limit chance peak matches. Peptide identification involved detecting and assembling the mass and intensity of the peptide peaks in MS spectra into three-dimensional peak hills over the m/z retention time plane. These were then filtered using graph theory algorithms to identify isotope patterns. Mass accuracy was achieved through weighted averaging and mass recalibration by subtracting systematic mass error from measured masses of each MS isotope pattern. Peptide and fragment masses (in the case of MS/MS spectra) were searched against an organism-specific sequence database. They were scored using a probability-based method called peptide score. The subsequent step involved assembling peptide hits into protein hits to identify proteins, where each identified peptide contributed to overall identification accuracy. The organism-specific database search included target sequences and their reverse counterparts and contaminants, aiding in determining a statistical cutoff for acceptable spectral matches. The main parameters set are shown in Table S3.

## Data analysis

All data were analyzed and plotted using the R package or GraphPad Prism software.

## Acknowledgments

The work was primarily supported by a research fund provided by the Science and Technology Development Program of Jilin Province, China (20220101062JC).

## Author Contributions

F.L. and L.P.L. designed and supervised the study. Y.W. and G.L. carried out experiments. Y.L., Y.P., and F.L. acquired and analyzed the data. Y.L. and F.L. drafted the manuscript. F.L. and L.P.L. revised the manuscript.

