## Supplemental Data for "Unveiling the Secrets of Extracellular Vesicles in Urban Water Systems: Understanding the Link Between Human and Environmental Health"

*Corresponding author:

The water sample of Guanlan River was characterized by measuring the following parameters in situ (Tabel S1):

Table S1 The Water Quality Data

| Sample | Time | Humidity | Temperature (℃) | Water Temperature (℃) | pH |
|---|---|---|---|---|---|
| S1 | 2022.12.03 5:04:00 PM | 95.3 | 19.7 | 22 | 7.37 |
| S2 | 2022.12.03 5:23:00 PM | 95.6 | 16.4 | 22.1 | 7.83 |
| S3 | 2022.12.03 5:30:00 PM | 96.4 | 17.8 | 23 | 8.04 |
| S4 | 2022.12.03 5:42:00 PM | 94.2 | 17.4 | 22.7 | 7.85 |
| S5 | 2022.12.03 5:54:00 PM | 92 | 16.6 | 20 | 8 |
| S6 | 2022.12.03 6:00:00 PM | 90.8 | 16.6 | 20.5 | 8 |

After a 1 μm prefiltration step, the water sample was isolated based on the EXODUS method[1] under an alternative vacuum actuation with the experimental condition of −20 kPa and 10-s conversion time. Following isolation, each subfraction was further eluted twice with PBS via EXODUS. Last, the reserved three fractions were recovered in 200 μl of 1X PBS and stored at −80°C for subsequent analysis. The details are shown in Table 2.

### Tabe S2. The Water Sample Process Procedure

| | | | | | | | | |
|---|---|---|---|---|---|---|---|---|
| **Separation condition** | 47mm, 500kd 50kPa, 10s 5 sonicate: High PBS 8mL*2 | | | 47mm, 500kd 50kPa, 10s 5 sonicate: High PBS 8mL*2 | 47mm, 500kd 50kPa, 10s 5 sonicate: High PBS 8mL*2 | 47mm, 500kd 50kPa, 10s 5 sonicate: High PBS 8mL*2 | 47mm, 500kd 50kPa, 10s 5 sonicate: High PBS 8mL*2 | 47mm, 500kd 50kPa, 10s 5 sonicate: High PBS 8mL*2 |
| **Particles recovery rate** | 11.58% | | | 11.28% | 13.14% | 13.54% | 4.39% | 8.67% |
| **Particle number/protein (particles/ug)** | 2.02E+09 | 6.17E+08 | 2.27E+08 | 1.35E+09 | 7.60E+08 | 1.45E+09 | 9.67E+08 | 1.50E+09 |
| **EVs mean size \(mean size, nm)** | 137.1 | 141.8 | 143.6 | 158.2 | 129.7 | 135.8 | 150.6 | 131.6 |
| **Average Size (nm)** | 151.6 | 151.6 | 151.6 | 170.2 | 153.4 | 161.8 | 167.1 | 163.6 |
| **EVs Particle Concentration (particles/mL)** | 2.2E+11 | 3.8E+10 | 1.2E+10 | 1.1 E+11 | 1.3E+11 | 1.3E+11 | 5.8E+10 | 1.3E+11 |
| **Sample NTA particle concentration (particles/mL)** | 3.8E+08 | 3.8E+08 | 3.8E+08 | 3.9E+08 | 3.1E+08 | 3.2E+08 | 5.9E+08 | 4.0E+08 |
| **EVs Protein (ug)** | 21.8 | 0 | 0 | 48.72 | 90.37 | 44.80 | 40.20 | 34.72 |
| **EVs Protein concentration (ug/mL)** | 109 | 62 | 53 | 81 | 171 | 90 | 60 | 87 |
| **Total Protein concentration (ug/mL)** | 22 | 22 | 22 | <10 | <10 | <10 | <10 | <10 |
| **EVs volume (μL)** | 200 | | | 600 | 470 | 500 | 670 | 400 |
| **Processing Time (min)** | 102 | | | 174 | 180 | 118 | 154 | 93 |
| **Processing Volume (mL)** | 1000 | | | 1500 | 1500 | 1500 | 1500 | 1500 |
| **Dilution** | 1 | | | 1 | 1 | 1 | 1 | 1 |
| **Volume (mL)** | 1000 | | | 1500 | 1500 | 1500 | 1500 | 1500 |
| **Sample #** | S1-2 | S1-2-1 | S1-2-2 | S2-1 | S3-1 | S4-1 | S5-1 | S6-1 |
| **Type** | Urban River | | | Urban River | Urban River | Urban River | Urban River | Urban River |
| **Equipment Info** | 31# | | | 31# | 31# | 31# | 31# | 31# |
| **Date** | 20221205 | | | 20221206 | 20221206 | 20221206 | 20221206 | 20221206 |
| **Note** | After the 1000ml sample is recovered, it is automatically emptied through the A10 chip and concentrated to a 200 ul final volume | Second Wash A01 Chip Result | Third Wash A01 Chip Result | | | | | |

**LC-MS/MS analysis**

All analyses were performed by a Q-Exactive HF mass spectrometer (Thermo, USA) equipped with a Nanospray Flex source (Thermo, USA). Samples were loaded and separated by a C18 column (25 cm × 75 μm) on an EASY-nLCTM 1200 system (Thermo, USA). The flow rate was 300nL/min and linear gradient was 90 min (0~1 min, 0%-2% B; 1~2 min, 2%-6% B; 2~51 min, 6%-21% B; 51~70 min, 21%-31% B; 70~81 min, 31%-43% B; 81~84 min, 43%-100% B; 84~90 min, 100%B; mobile phase A = 0.1% FA in water and B = 0.1% FA in 80% CAN and 19.9% water). Full MS scans were acquired in the mass range of 350-1650 m/z with a mass resolution of 60000, and the AGC target value was set at 3e6. The 20 most intense peaks in MS were fragmented with higher-energy collisional dissociation (HCD) with a collision energy of 28. MS/MS spectra were obtained with a resolution of 30000 with an AGC target of 2e5 and a max injection time of 80 ms. The Q Exactive HF dynamic exclusion was set for 40.0 s and run under positive mode.

**Database search**

The LC-MS/MS raw data were imported in Maxquant (Version 1.6.17.0) for labeling-free quantification analysis, and the search engine was Andromeda. A target-decoy-based false discovery rate (FDR) approach limits a certain number of peak matches by chance. For peptide identification, the mass and intensity of the peptide peaks in mass spectrometry (MS) spectra are detected and assembled into three-dimensional (3D) peak hills over the m/z retention time plane, which is filtered by applying graph theory algorithms to identify isotope patterns. High mass accuracy is achieved by weighted averaging and through mass recalibration by subtracting the determined systematic mass error from the measured mass of each MS isotope pattern. Peptide and fragment masses (in the case of an MS/MS spectra) are searched in an organism-specific sequence database and are then scored by a probability-based approach termed peptide score. The next step is assembling peptide hits into protein hits to identify proteins. The assembly of peptide hits into protein hits to identify proteins is the next step, in which each identified peptide of a protein contributes to the overall identification accuracy. The organism-specific database search includes the target sequences and their reverse counterparts and contaminants, which helps to determine a statistical cutoff for acceptable spectral matches. The main parameters were set as follows (Table S3):

| Items | Settings |
|---|---|
| FDR | 0.01 |
| Missed cleavage | 2 |
| Fixed modification | Carbamidomethyl（C） |
| Variable modification | Oxidation（M）、Acetyl（Protein N-term） |
| Decoy database pattern | Reverse |
| enzyme | Trypsin |
| First search for peptide tolerance | 20 ppm |
| Main search peptide tolerance | 4.5 ppm |
| Database | |

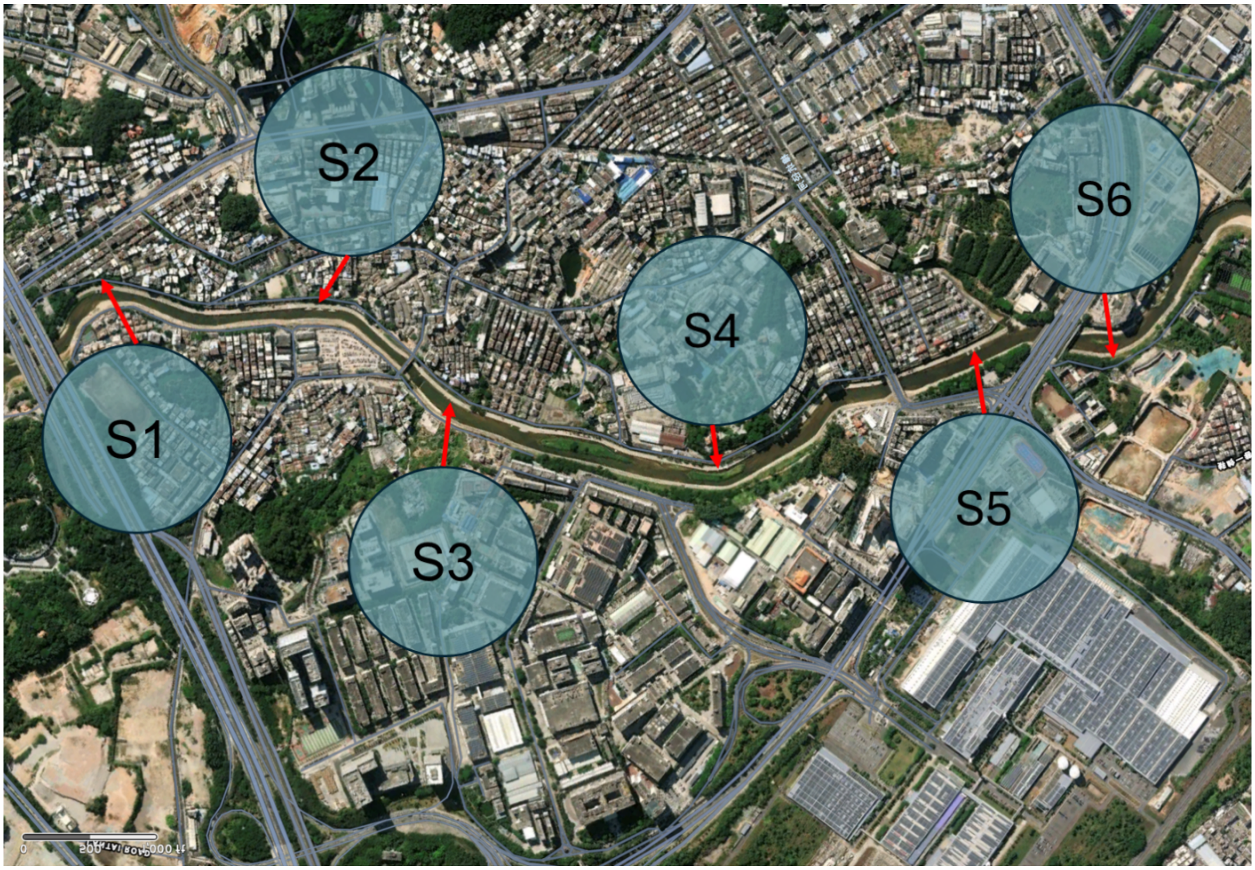


Fig. S1 The samples were collected from 6 different locations along Guanlan River.

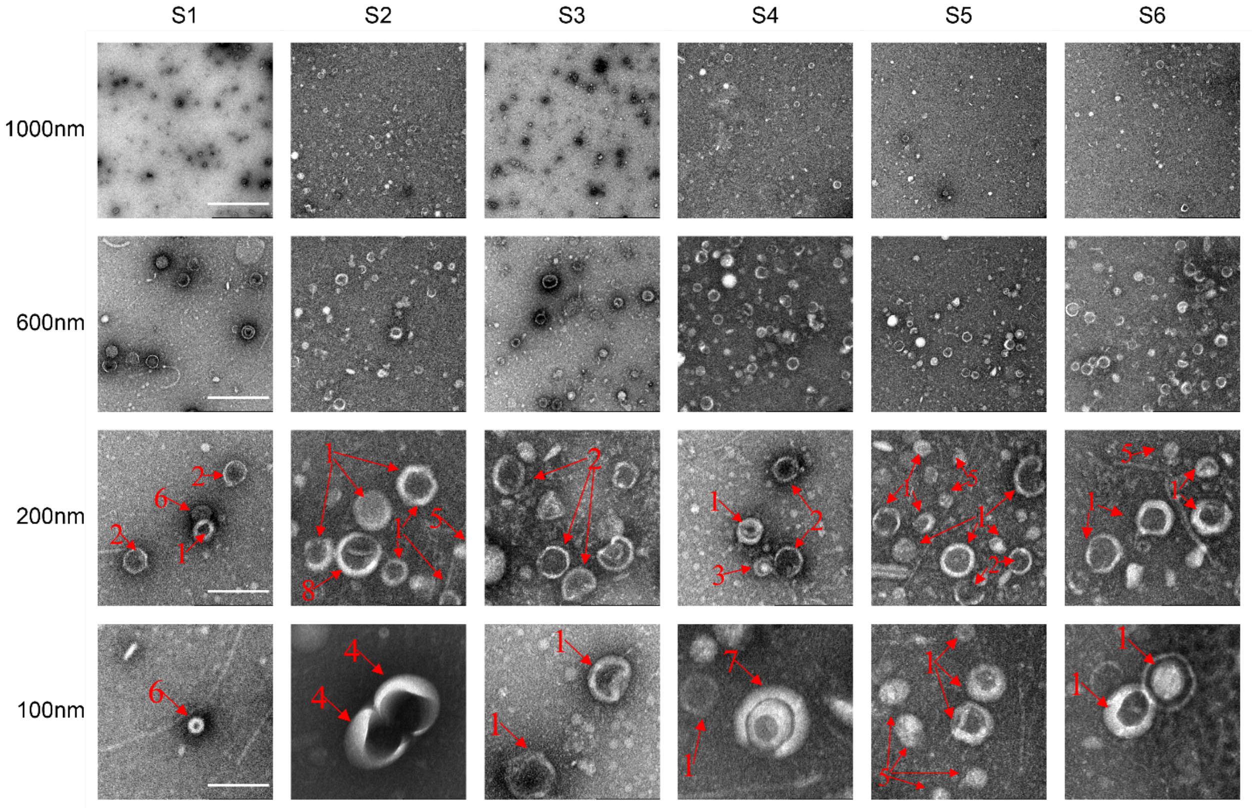


Fig. S2 Various magnification levels were utilized to capture TEM images of samples S1 through S6, providing comprehensive visual portrayals of the morphology, structure, and distinguishing attributes of the standard EVs at varying scales or resolutions within each sample. First-row scale bar: 1000nm, second-row scale bar: 400nm, third-row scale bar: 200nm, fourth-row scale bar: 100nm.

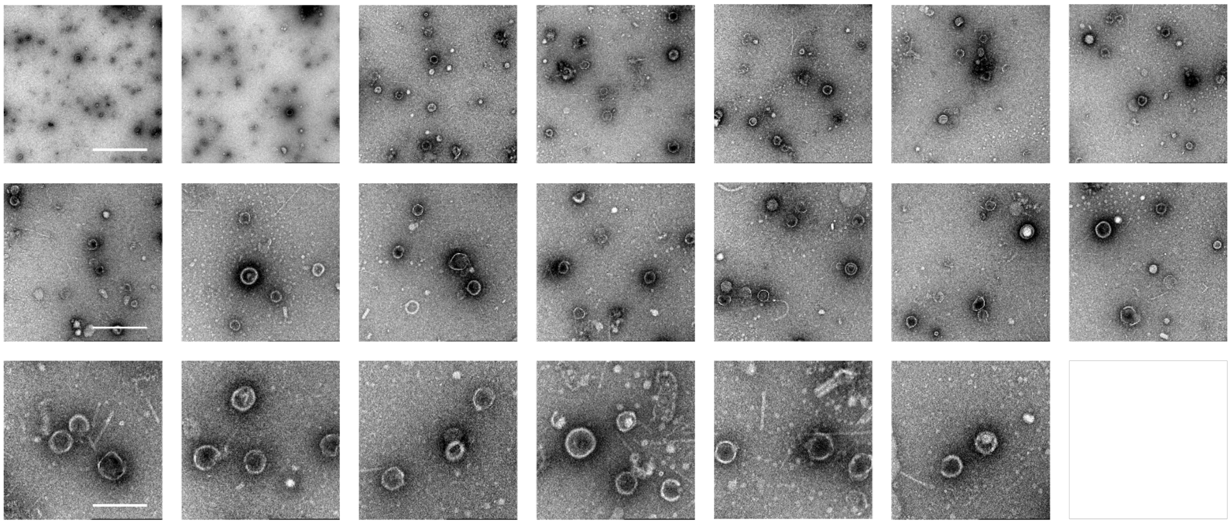

Fig. S3 shows various magnification levels utilized to capture TEM images of Sample S1, offering a detailed visual representation of the EV's morphology, structure, and characteristics at different scales or resolutions. The first-row scale bar is 1000nm, the second-row scale bar is 600nm, and the third-row scale bar is 200nm.

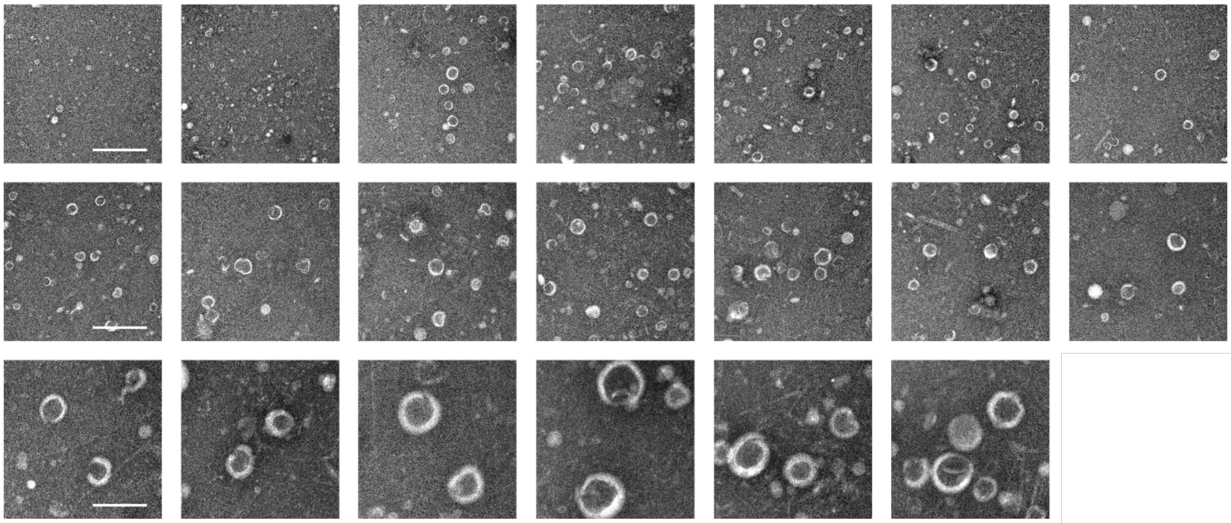

Fig. S4 Various magnification levels were utilized to capture TEM images of Sample S2, offering a detailed visual representation of the EV's morphology, structure, and characteristics at different scales or resolutions. The first-row scale bar is 1000nm, the second-row scale bar is 600nm, and the third-row scale bar is 200nm.

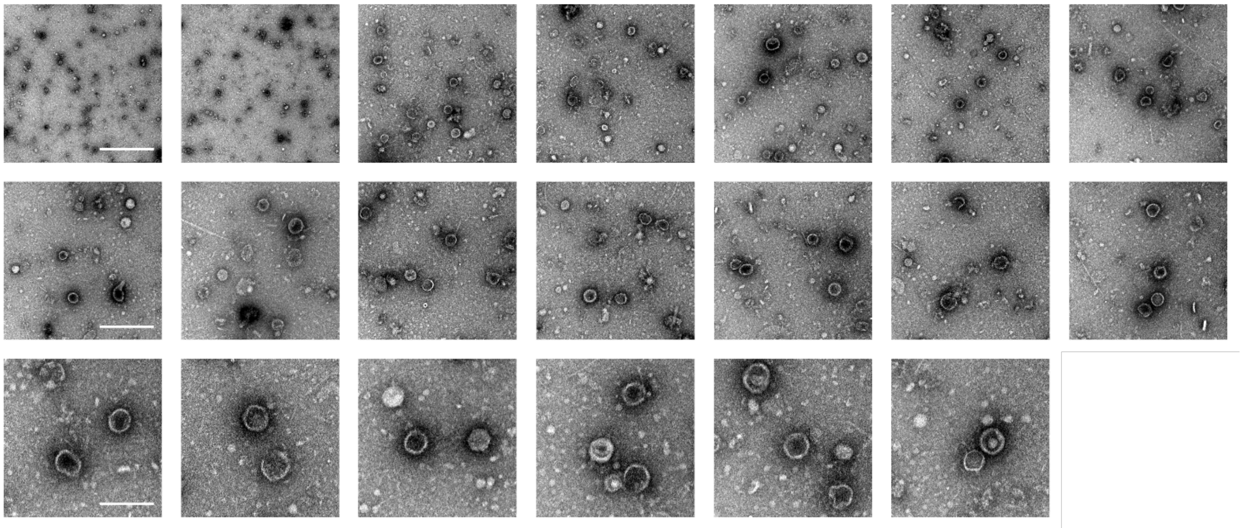

Fig. S5 Various magnification levels were utilized to capture TEM images of Sample S3, offering a detailed visual representation of the EV's morphology, structure, and characteristics at different scales or resolutions. The first-row scale bar is 1000nm, the second-row scale bar is 400nm, and the third-row scale bar is 200nm.

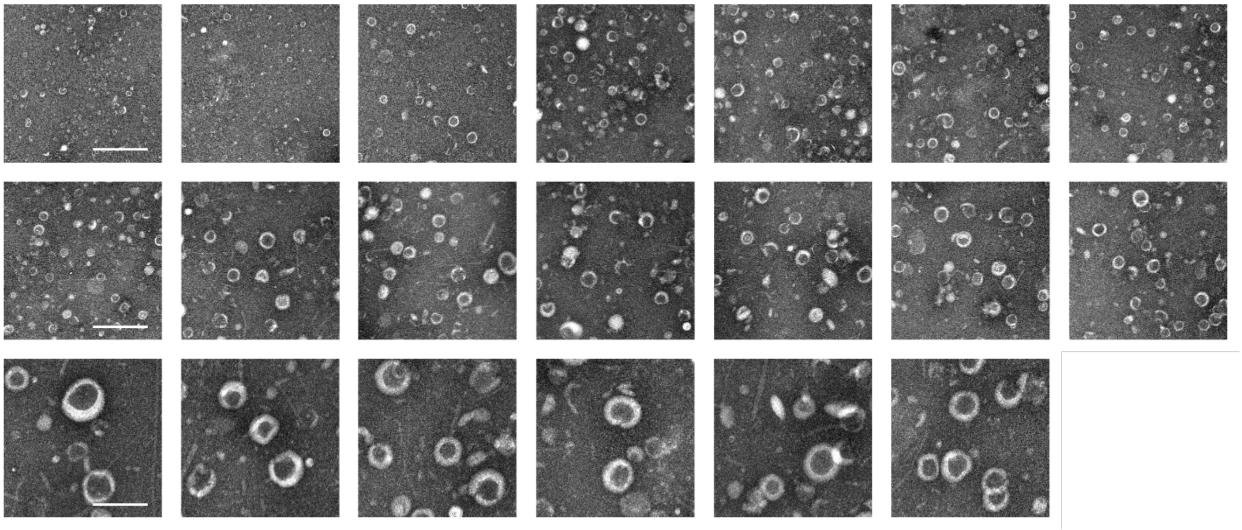

Fig. S6 Various magnification levels were utilized to capture TEM images of Sample S4, offering a detailed visual representation of the EV's morphology, structure, and characteristics at different scales or resolutions. The first-row scale bar is 1000nm, the second-row scale bar is 400nm, and the third-row scale bar is 200nm.

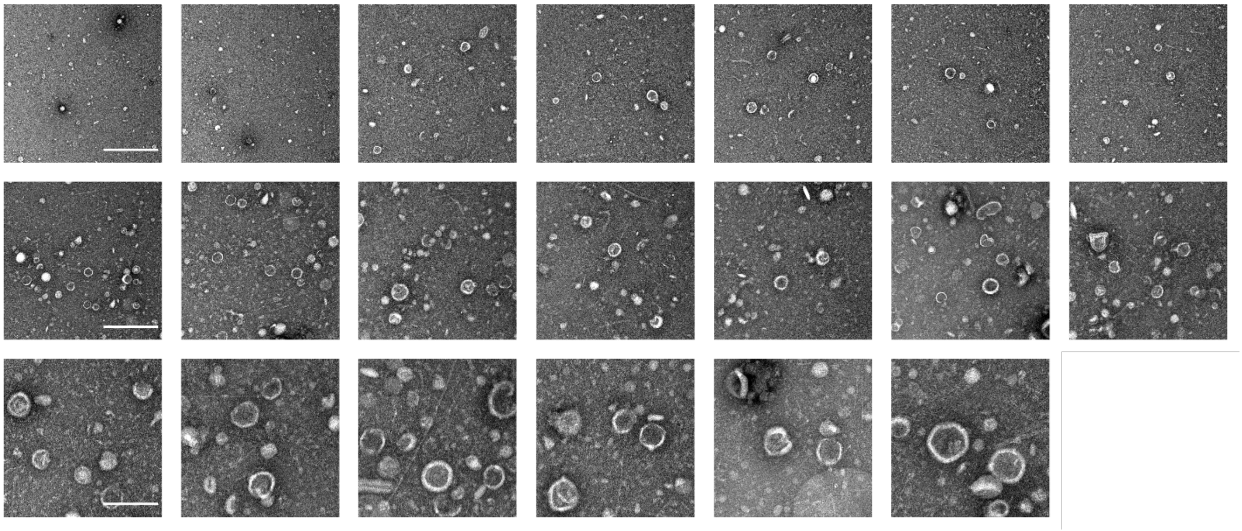

Fig. S7 Various magnification levels were utilized to capture TEM images of Sample S5, offering a detailed visual representation of the EV's morphology, structure, and characteristics at different scales or resolutions. The first-row scale bar is 1000nm, the second-row scale bar is 400nm, and the third is 200nm.

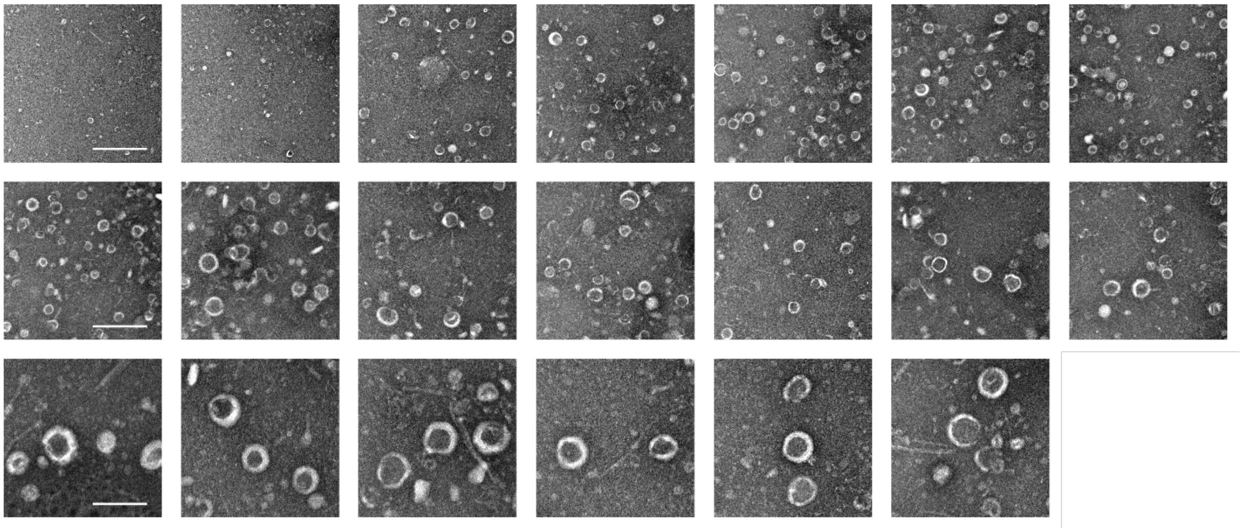

Fig. S8 Various magnification levels were utilized to capture TEM images of Sample S6, offering a detailed visual representation of the EV's morphology, structure, and characteristics at different scales or resolutions. The first-row scale bar is 1000nm, the second-row scale bar is 400nm, and the third-row scale bar is 200nm.

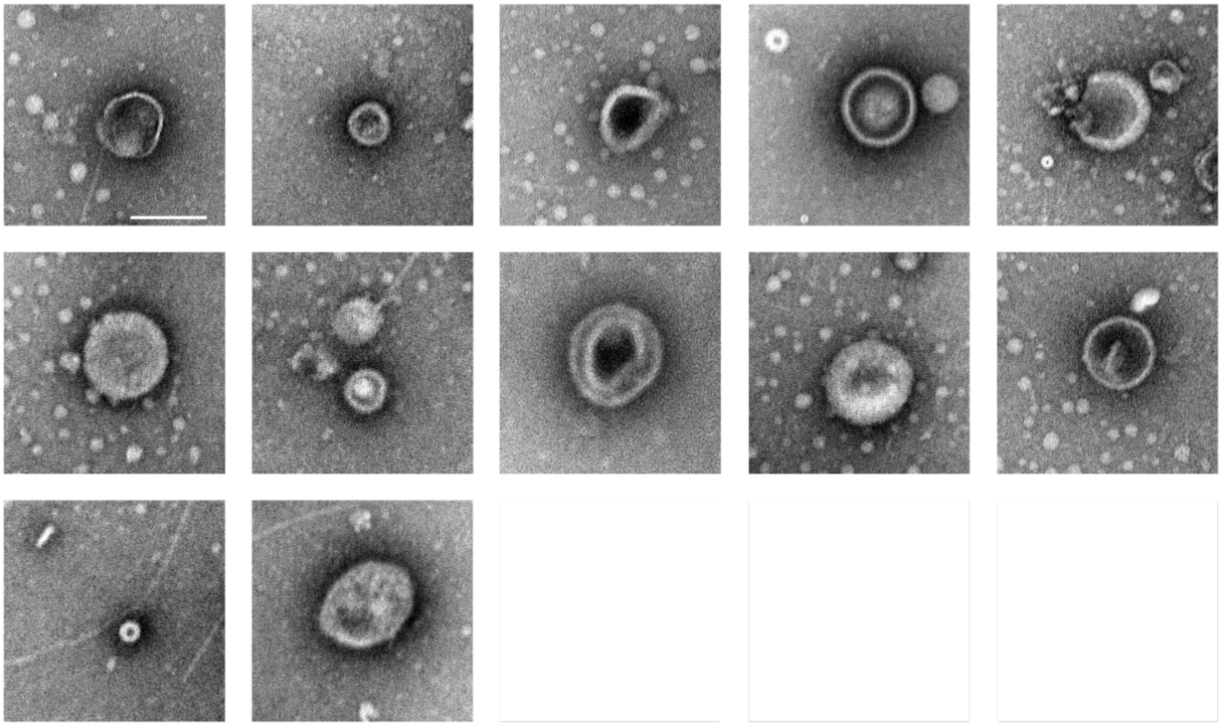

Fig. S9 TEM images of sample S1 present a highly detailed and precise visual rendering, explicitly showcasing the intricate morphology, structural components, and unique characteristics of individual EVs within the sample. Scale bar: 100nm.

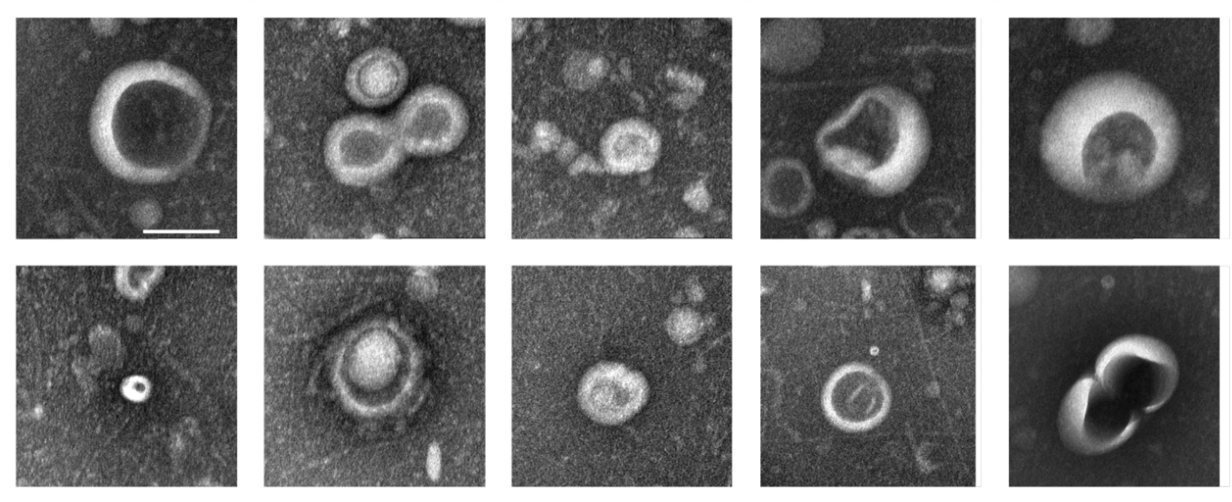

Fig. S10 TEM images of sample S2 present a highly detailed and precise visual rendering, explicitly showcasing the intricate morphology, structural components, and unique characteristics of individual EVs within the sample. Scale bar: 100nm.

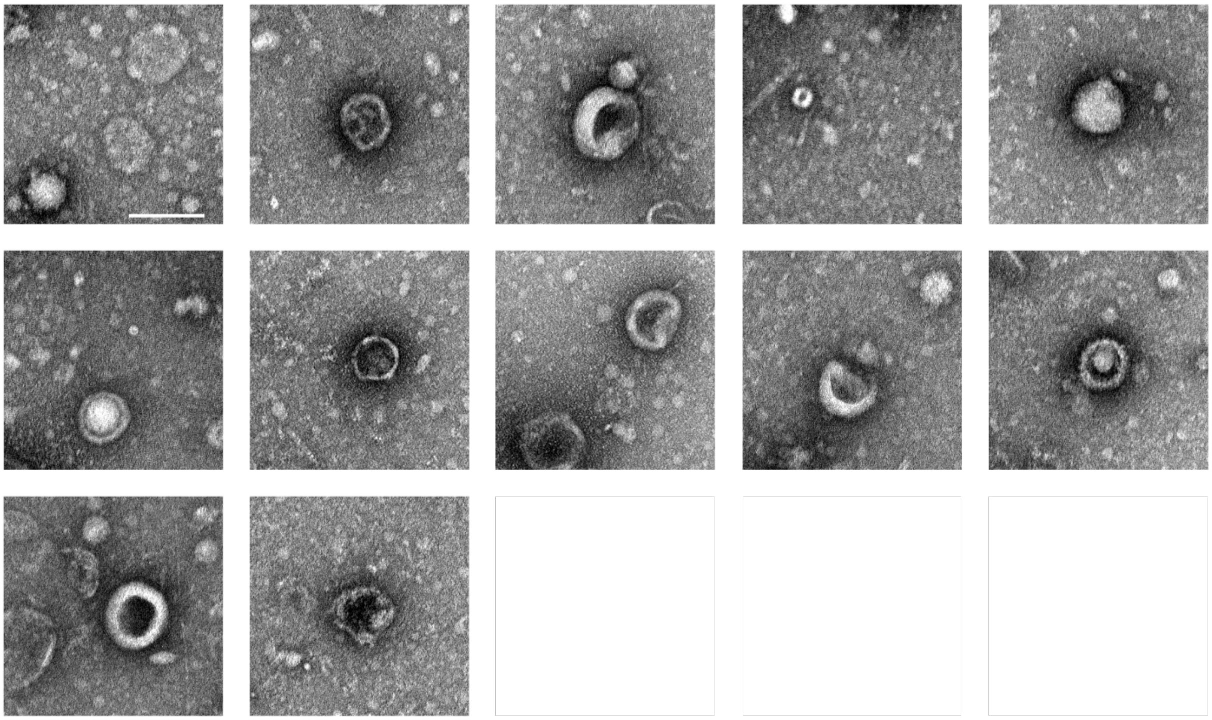

Fig. S11 TEM images of sample S3 present a highly detailed and precise visual rendering, explicitly showcasing the intricate morphology, structural components, and unique characteristics of individual EVs within the sample. Scale bar: 100nm.

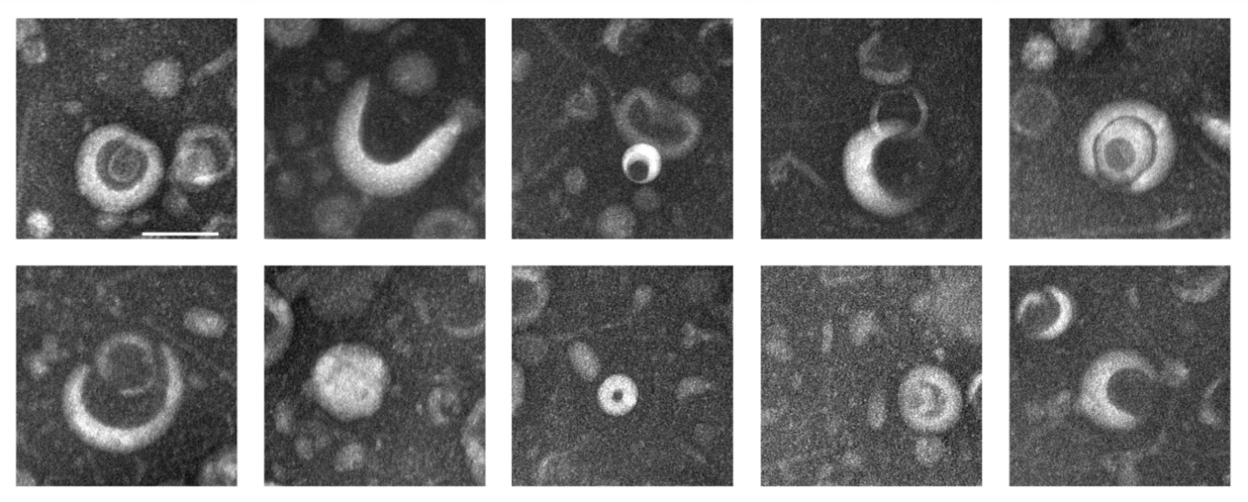

Fig. S12 TEM images of Sample S4s present a highly detailed and precise visual rendering, explicitly showcasing the intricate morphology, structural components, and unique characteristics of individual EVs within the sample. Scale bar: 100nm.

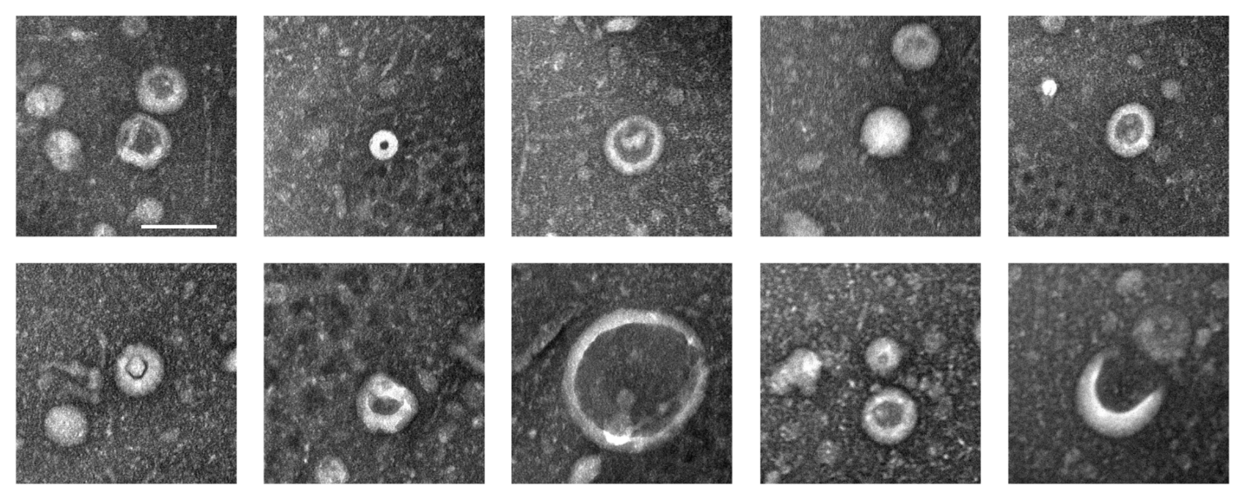

Fig. S13 TEM images of sample S5 present a highly detailed and precise visual rendering, explicitly showcasing the intricate morphology, structural components, and unique characteristics of individual EVs within the sample. Scale bar: 100nm.

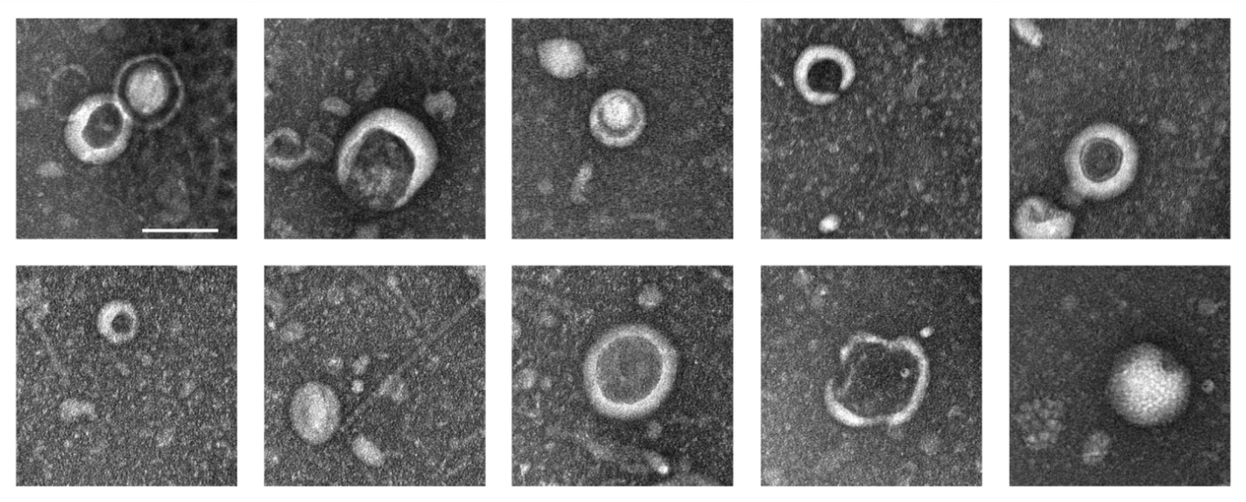

Fig. S14 TEM images of sample S6 present a highly detailed and precise visual rendering, explicitly showcasing the intricate morphology, structural components, and unique characteristics of individual EVs within the sample. Scale bar: 100nm.

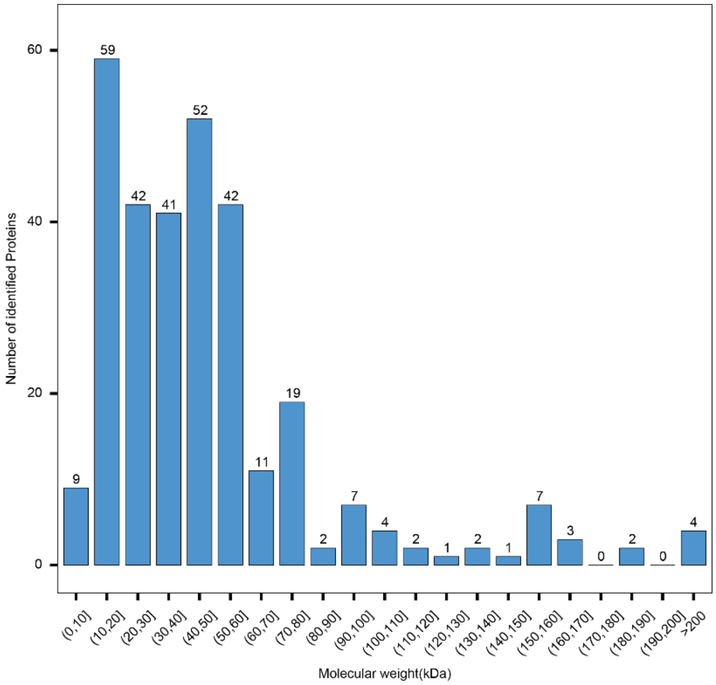


Fig. S15 The distribution of molecular weights observed across 6 different EV samples.

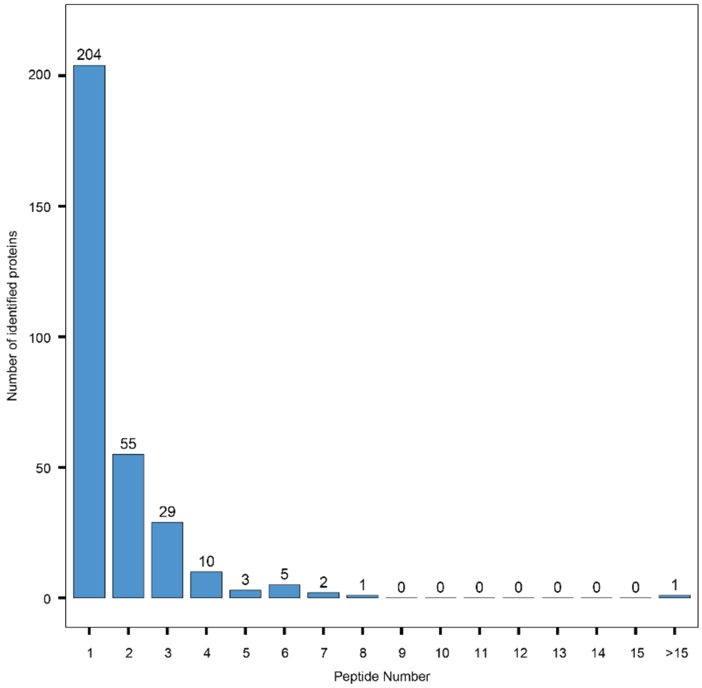


Fig. S16 Peptide number distribution observed across 6 different EV samples.

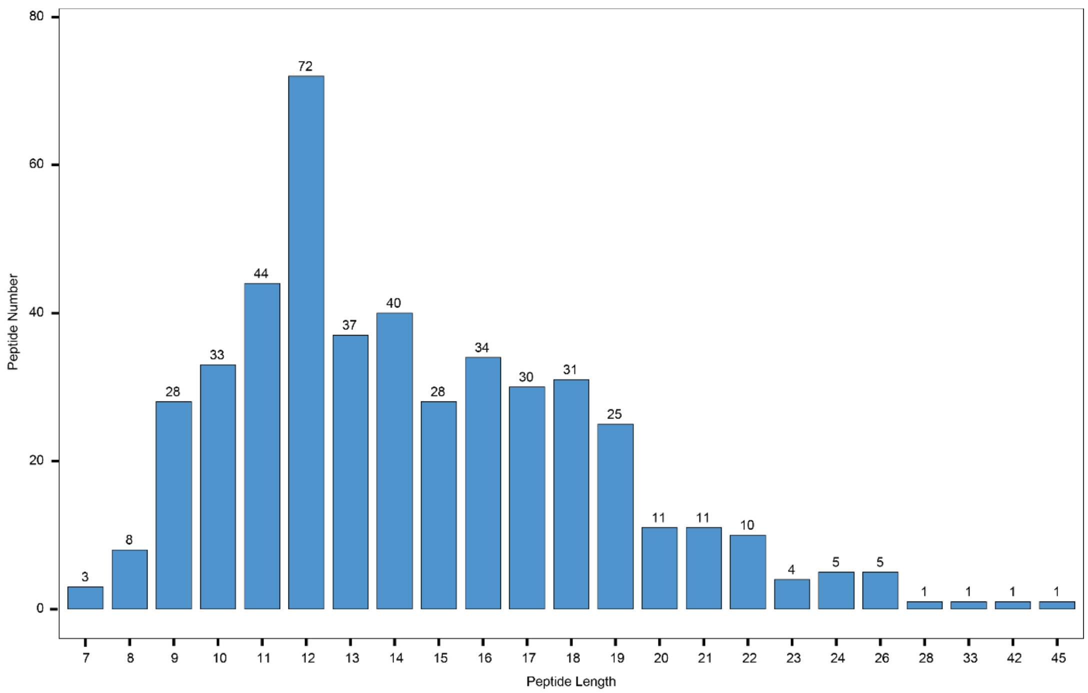


Fig. S17 Peptide length distribution observed across 6 different EV samples.

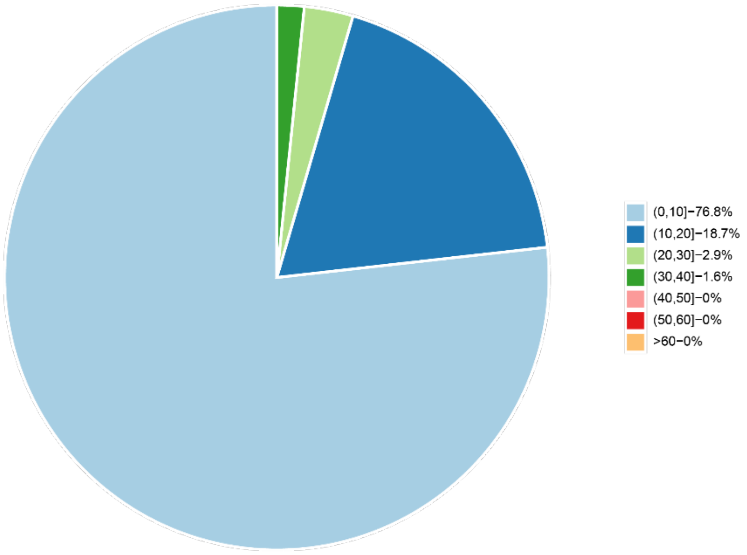


Fig. S18 Peptide coverage observed across the entirety of 6 different EV samples.

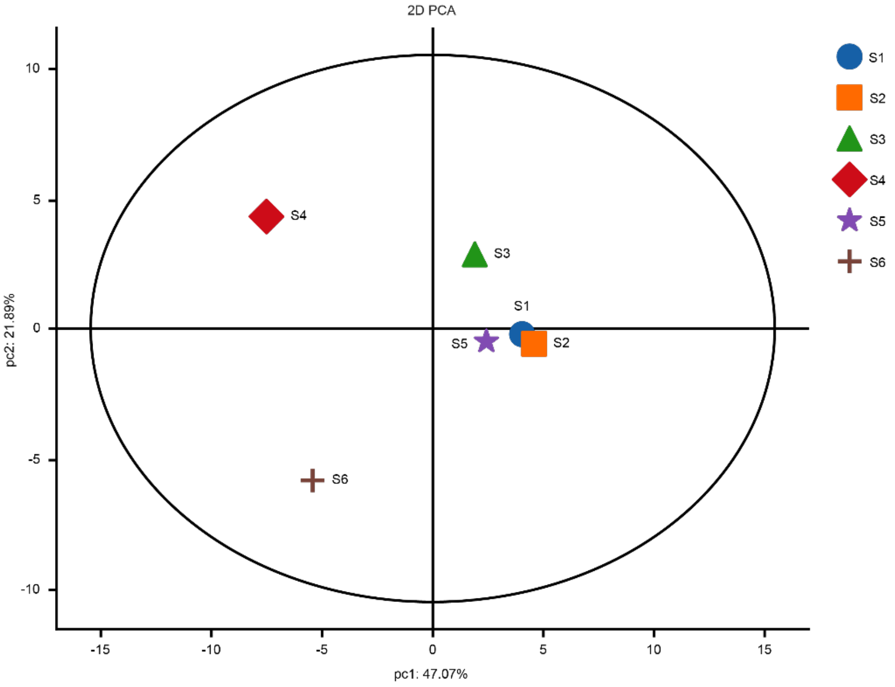


Fig. S19 PCA conducted in a 2D format encompassing all 6 distinct EV samples.

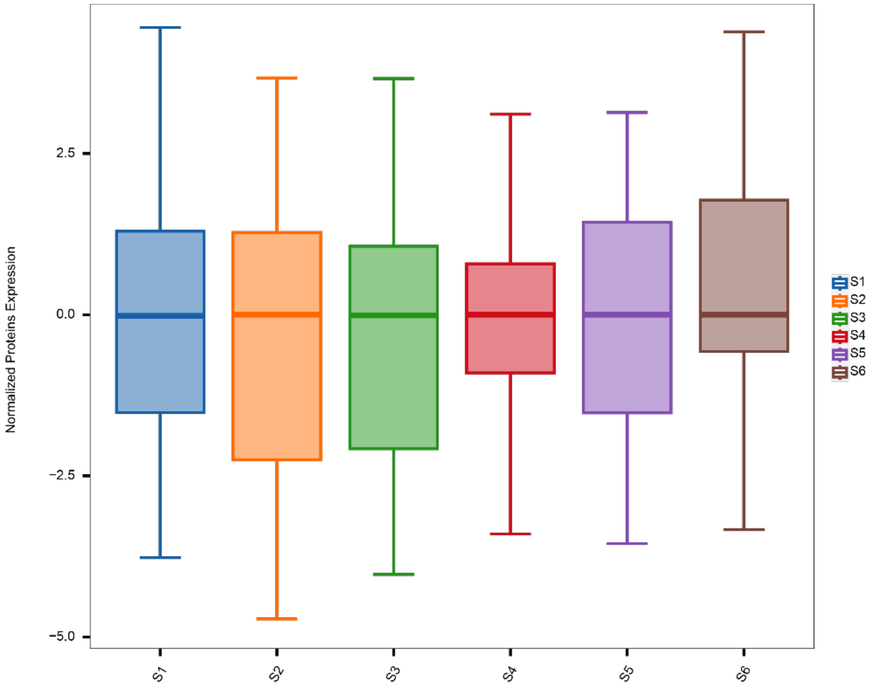

Fig. S20 Normalized protein expression level in each of the 6 EV samples in total.

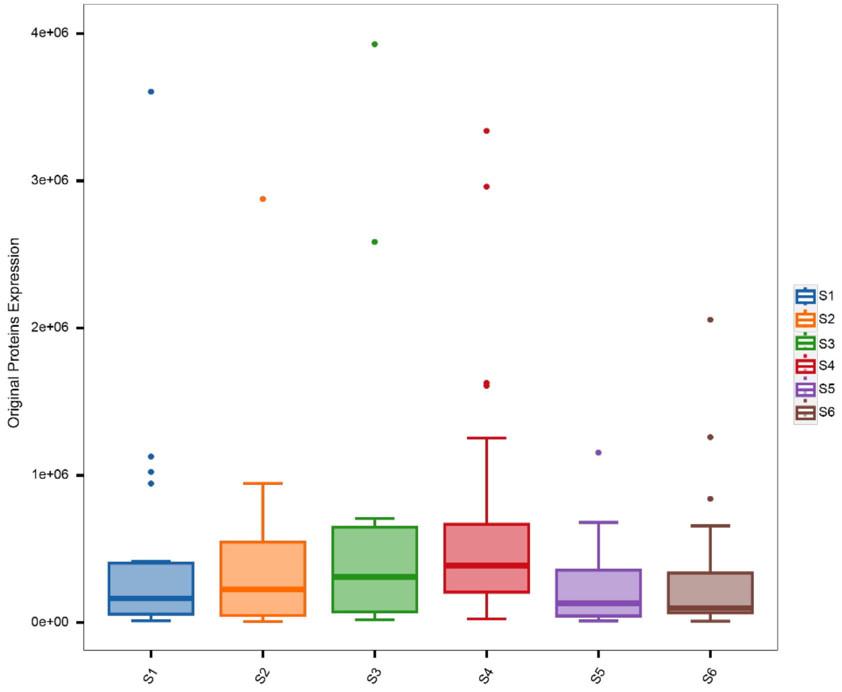


Fig. S21 Original protein expression level in each of the 6 EV samples in total.

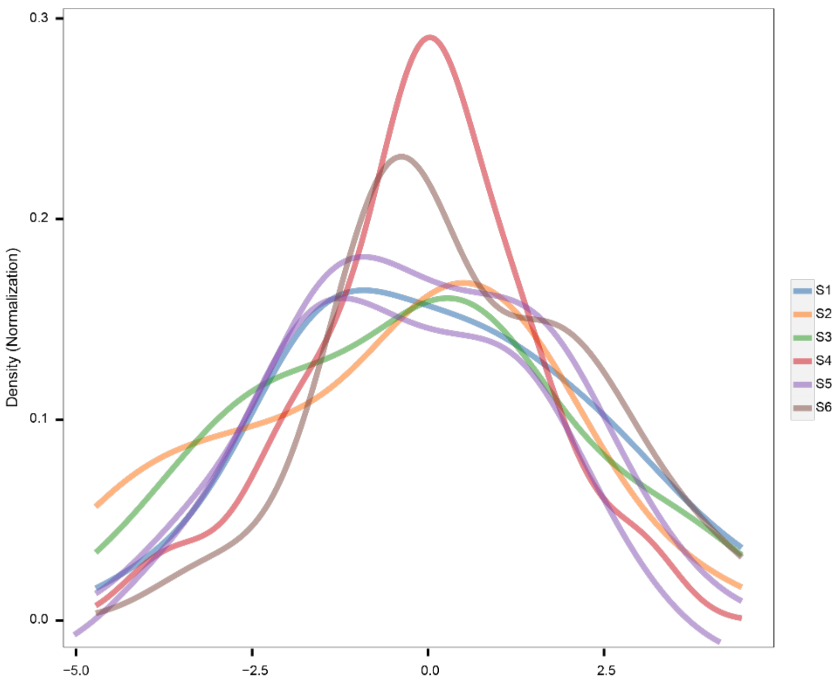


Fig. S22 Normalized protein expression density in each of the 6 EV samples in total.

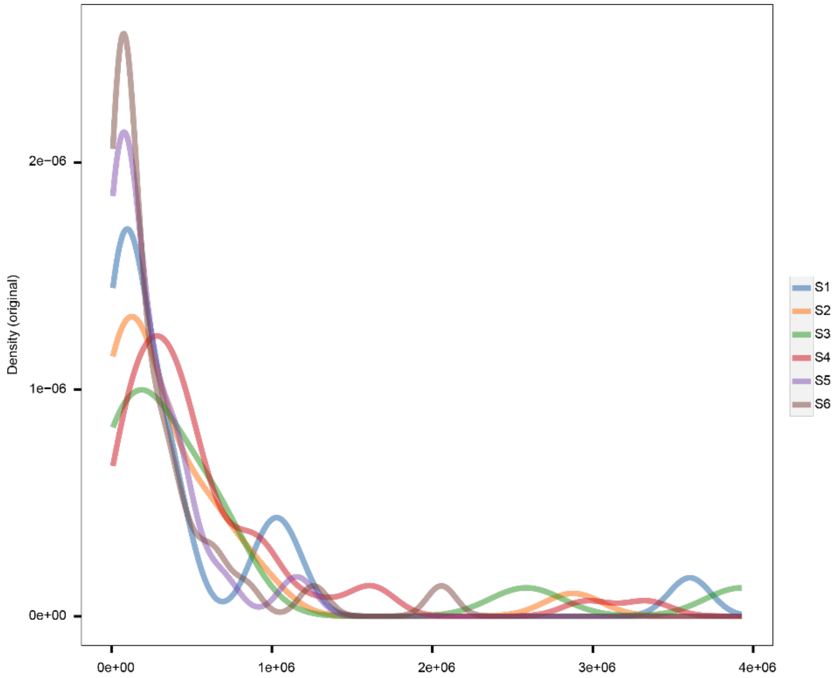


Fig. S23 Original protein expression density in each of the 6 EV samples in total.

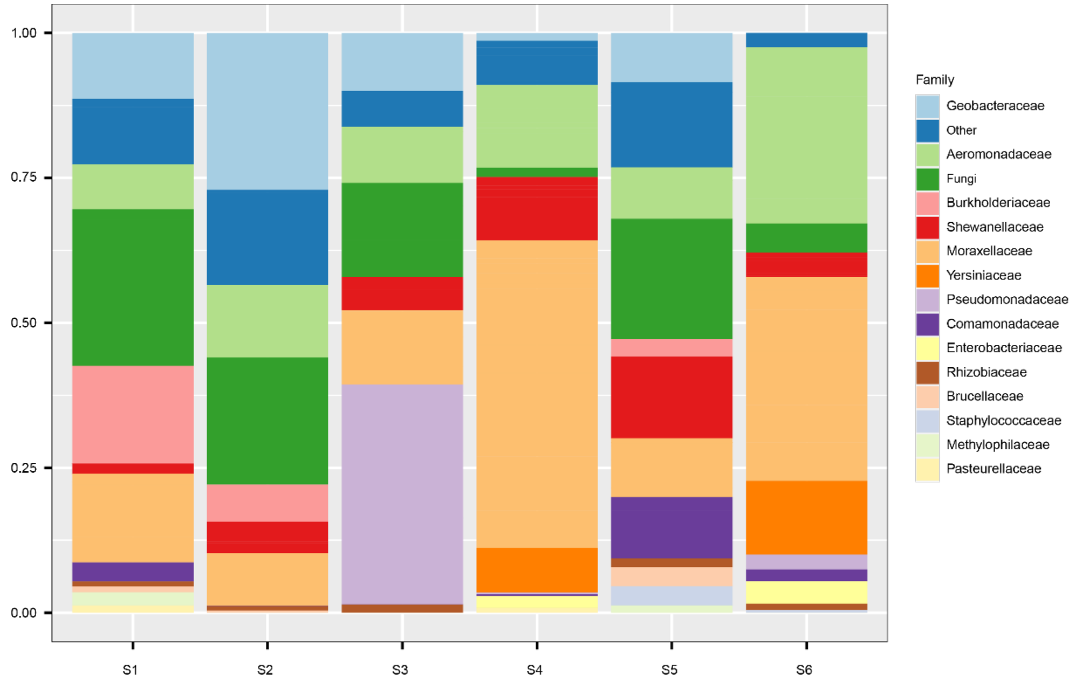


Fig. S24 Bacterial family distribution in each of the 6 EV samples in total.

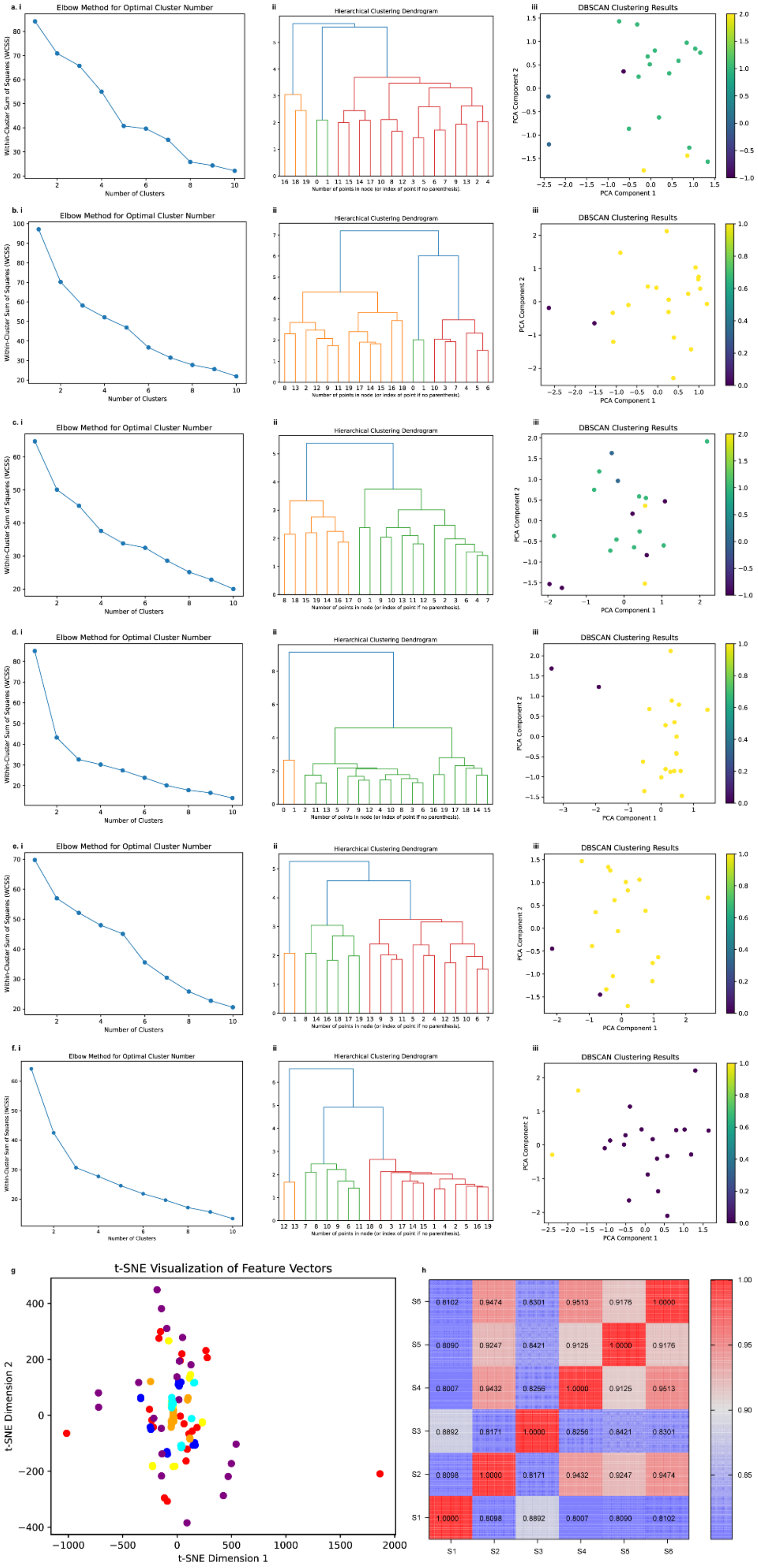
a. i
Elbow Method for Optimal Cluster Number
Within-Cluster Sum of Squares (WCSS)
Number of Clusters
ii
Hierarchical Clustering Dendrogram
Number of points in node (or index of point if no parenthesis).
iii
DBSCAN Clustering Results
PCA Component 1
PCA Component 2
b. i
c. i
d. i
e. i
f. i
g
t-SNE Visualization of Feature Vectors
t-SNE Dimension 1
t-SNE Dimension 2
h
S6 0.8102 0.9474 0.8301 0.9513 0.9176 1.0000
S5 0.8090 0.9247 0.8421 0.9125 1.0000 0.9176
S4 0.8007 0.9432 0.8256 1.0000 0.9125 0.9513
S3 0.8892 0.8171 1.0000 0.8256 0.8421 0.8301
S2 0.8098 1.0000 0.8171 0.9432 0.9247 0.9474
S1 1.0000 0.8098 0.8892 0.8007 0.8090 0.8102
S1 S2 S3 S4 S5 S6

Fig. S25 The ViT clustering results. **a (i) - f (i)** The Elbow method, used to cluster images within each image set, involves assessing the within-cluster sum of squares (WCSS) alongside computing the silhouette coefficient. This combined approach identifies the optimal number of clusters (represented as K in the K-means clustering method). **a (ii) - f (ii)** A dendrogram for hierarchical clustering is utilized to cluster images within each set. This method offers detailed understanding at each stage of merging clusters, producing a dendrogram in the process. **a (iii) - f (iii)** density-based spatial clustering of applications with noise (DBSCAN) clustering offers a more intricate analysis of clustering for each set of images based on density. **g** the feature vectors extracted from each collection of images underwent clustering using the T-distributed stochastic neighbor embedding (t-SNE) technique. **h** the image feature vectors are analyzed for correlation using the Structural Similarity Index Measure (SSIM) technique. Findings show that EVs originating from S4 and S6 demonstrate the closest resemblance in their morphology.

Table S3 Top three Bacteria’s family in sample S1 – S6

| S1 | S2 | S3 | S4 | S5 | S6 |
|---|---|---|---|---|---|
| Fungi | ***Geobacteraceae*** | ***Pseudomonadaceae*** | ***Moraxellaceae*** | Fungi | ***Moraxellaceae*** |
| ***Burkholderiaceae*** | Fungi | Fungi | *Aeromonadaceae* | other | *Aeromonadaceae* |
| *Moraxellaceae* | other | *Geobacteraceae* | *Shewanellaceae* | ***Shewanellaceae*** | *Yersiniaceae* |

1 Chen, Y. *et al.* Exosome detection via the ultrafast-isolation system: EXODUS. *Nature Methods* **18**, 212-218 (2021).
